# Sustained software development, not number of citations or journal choice, is indicative of accurate bioinformatic software

**DOI:** 10.1101/092205

**Authors:** Paul P. Gardner, James M. Paterson, Stephanie McGimpsey, Fatemeh Ashari-Ghomi, Sinan U. Umu, Aleksandra Pawlik, Alex Gavryushkin, Michael A. Black

## Abstract

**Background:** Computational biology provides widely used and powerful software tools for testing and making inferences about biological data. In the face of rapidly increasing volumes of data, heuristic methods that trade software speed for accuracy may be employed. We are have studied these trade-offs using the results of a large number of independent software benchmarks, and evaluated whether external factors are indicative of accurate software.

**Method:** We have extracted accuracy and speed ranks from independent benchmarks of different bioinformatic software tools, and evaluated whether the speed, author reputation, journal impact, recency and developer efforts are indicative of accuracy.

**Results:** We found that software speed, author reputation, journal impact, number of citations and age are all unreliable predictors of software accuracy. This is unfortunate because citations, author and journal reputation are frequently cited reasons for selecting software tools. However, GitHub-derived records and high version numbers show that the accurate bioinformatic software tools are generally the product of many improvements over time, often from multiple developers.

**Discussion:** We also find that the field of bioinformatics has a large excess of slow and inaccurate software tools, and this is consistent across many sub-disciplines. Meanwhile, there are few tools that are middle-of-road in terms of accuracy and speed trade-offs. We hypothesise that a form of publication-bias influences the publication and development of bioinformatic software. In other words, software that is intermediate in terms of both speed and accuracy may be difficult to publish - possibly due to author, editor and reviewer practices. This leaves an unfortunate hole in the literature as the ideal tools may fall into this gap. For example, high accuracy tools are not always useful if years of CPU time are required, while high speed is not useful if the results are also inaccurate.

## Background

Computational biology software is widely used and has produced some of the most cited publications in the entire scientific corpus [1, 2, 3]. These highly-cited software tools include implementations of methods for sequence alignment and homology inference [4, 5, 6, 7], phylogenetic analysis [8, 9, 10, 11, 12], biomolecular structure analysis [13, 14, 15, 16, 17], and visualization and data collection [18, 19]. However, the popularity of a software tool does not necessarily mean that it is accurate or computationally efficient; instead usability, ease of installation, operating system support or other indirect factors may play a greater role in a software tool’s popularity. Indeed, there have been several notable incidences where convenient, yet inaccurate software has caused considerable harm [20, 21, 22].

Progress in the biological sciences is increasingly limited by the ability to analyse large volumes of data, therefore the dependence of biologists on software is also increasing [23]. There is an increasing reliance on technological solutions for automating biological data generation (e.g. next-generation sequencing, mass-spectroscopy, cell-tracking and species tracking), therefore the biological sciences have become increasingly dependent upon software tools for processing large quantities of data [23]. As a consequence, the computational efficiency of data processing and analysis software is of great importance to decrease the energy, climate impact, and time costs of research [24]. Furthermore, as datasets become larger even small error rates can have major impacts on the number of false inferences [25].

The gold-standard for determining accuracy is for researchers independent of individual tool development to conduct benchmarking studies; these benchmarks can serve a useful role in reducing the over-optimistic reporting of software accuracy [26, 27, 28] and the self-assessment trap [29, 30]. Benchmarking typically involves the use a number of positive and negative control datasets, so that predictions from different software tools can be partitioned into true or false groups, allowing a variety of metrics to be used to evaluate performance [31, 32, 28]. The aim of these benchmarks is to robustly identify tools that make acceptable compromises in terms of balancing speed with discriminating true and false predictions, and are therefore suited for wide adoption by the community.

For common computational biology tasks, a proliferation of software-based solutions often exists [33, 34, 35]. While this is a good problem to have, and points to a diversity of options from which practical solutions can be selected, having many possible options creates a dilemma for users. In the absence of any recent gold-standard benchmarks, how should scientific software be selected? In the following we presume that the “biological accuracy” of predictions is the most desirable feature for a software tool. Biological accuracy is the degree to which predictions or measurements reflect the biological truths based on expert-derived curated datasets (see Methods for the mathematical definition used here).

A number of possible predictors of software quality are used by the community of computational biology software users [36, 37, 38]. Some accessible, quantifiable and frequently used proxies for identifying high quality software include: **1. Recency:** recently published software tools may have built upon the results of past work, or be an update to an existing tool. Therefore these may be more accurate and faster. **2. Wide adoption:** a software tool may be widely used because it is fast and accurate, or because it is well-supported and user-friendly. In fact,”large user base”, “word-of-mouth”, “wide-adoption”, “personal recommendation”, and “recommendation from a close colleague”, were frequent responses to surveys of “how do scientists select software?” [36, 37, 38]. **3. Journal impact:** many believe that high profile journals are run by editors and reviewers who carefully select and curate the best manuscripts. Therefore, high impact journals may be more likely to select manuscripts describing good software [39]. **4. Author/group reputation:** the key to any project is the skills of the people involved, including maintaining a high collective intelligence [37, 40, 41]. As a consequence, an argument could be made that well respected and high-profile authors may write better software [42, 43]. **5. Speed:** software tools frequently trade accuracy for speed. For example, heuristic software such as the popular homology search tool, BLAST, compromises the mathematical guarantee of optimal solutions for more speed [4, 7]. Some researchers may naively interpret this fact as implying that slower software is likely to be more accurate. But speed may also be influenced by the programming language [44], and the level of hardware optimisation [45, 46]; however, the specific method of implementation generally has a greater impact (e.g., brute-force approaches versus rapid and sensitive pre-filtering [47, 48, 49]). **6. Effective software versioning:** With the wide adoption of public version-control systems like GitHub, quantifiable data on software development time and intensity indicators, such as the number of contributors to code, number of code changes and versions is now available [50, 51, 52].

In the following study, we explore factors that may be indicative of software accuracy. This, in our opinion, should be one of the prime reasons for selecting a software tool. We have mined the large and freely accessible PubMed database [53] for benchmarks of computational biology software, and manually extracted accuracy and speed rankings for 499 unique software tools. For each tool, we have collected measures that may be predictive of accuracy, and may be subjectively employed by the research community as a proxy for software quality. These include relative speed, relative age, the productivity and impact of the corresponding authors, journal impact, number of citations and GitHub activity.

## Results

We have collected relative accuracy and speed ranks for 499 distinct software tools. This software has been developed for solving a broad cross-section of computational biology tasks. Each software tool was benchmarked in at least one of 69 publications that satisfy the Boulesteix criteria [54]. In brief, the Boulesteix criteria are: 1. the main focus of the article is a benchmark. 2. the authors are reasonably neutral. 3. the test data and evaluation criteria are sensible.

For each of the publications describing these tools, we have (where possible) collected the journal’s H5-index (Google Scholar Metrics), the maximum H-index and corresponding M-indices [42] for the corresponding authors for each tool, and the number of times the publication(s) associated with a tool has been cited using Google Scholar (data collected over a 6 month period in late 2020). Note that citation metrics are not static and will change over time. In addition, where possible we also extract the version number, the number of commits, number of contributors total number “issues”, the proportion of issues that remain open, the number of pull requests, and the number of times the code was forked from public GitHub repositories.

We have computed the Spearman’s correlation coefficient for each pairwise combination of the mean normalised accuracy and speed ranks, with the year published, mean relative age (compared to software in the same benchmarks), journal H5 metrics, the total number of citations, the relative number of citations (compared to software in the same benchmarks) and the maximum H- and corresponding M-indices for the corresponding authors, version number, and the GitHub statistics commits, contributors, pull requests, issues, % open issues and forks. The results are presented in Figure 1A&B, and Additional file 1: Figures S5&S6. We find significant associations between most of the citation-based metrics (journal H5, citations, relative citations, H-index and M-index). There is also a negative correlation between the year of publication, the relative age and many of the citation-based metrics.

**Figure 1:**
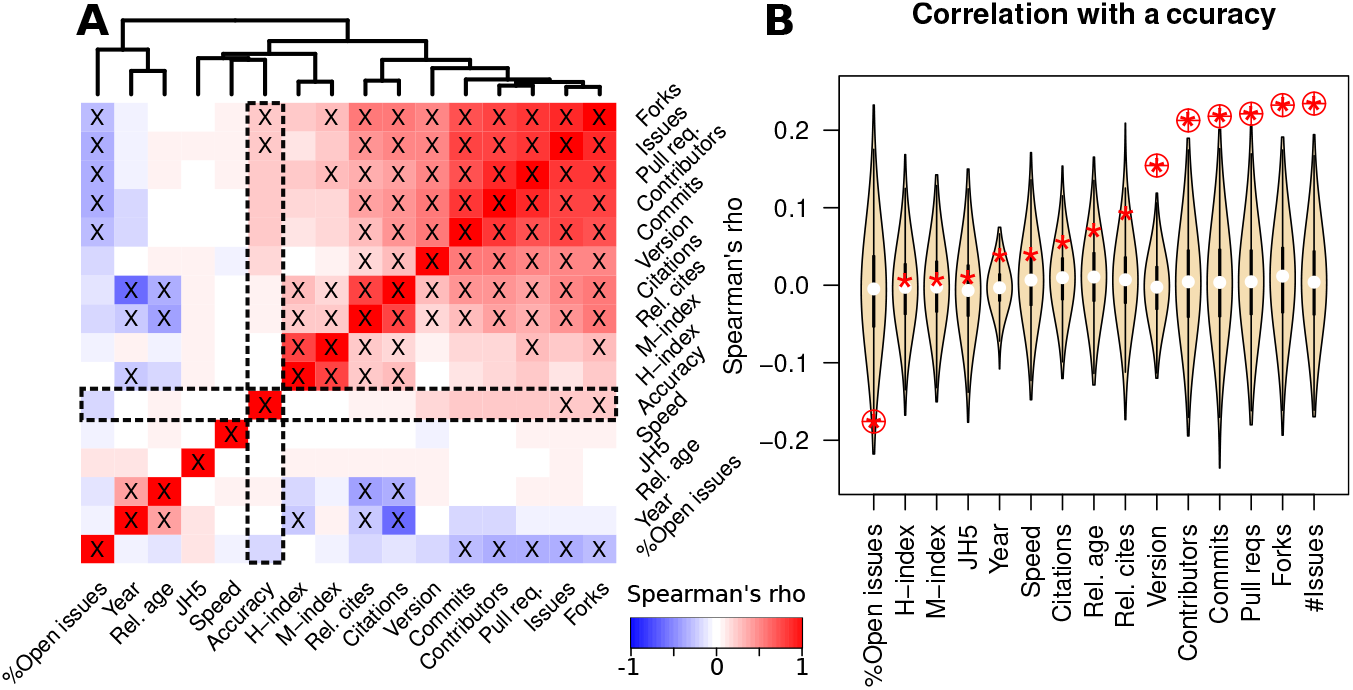
**A.** A heatmap indicating the relationships between different features of bioinformatic software tools. Spearman’s rho is used to infer correlations between metrics such as citations based metrics, the year and relative age of publication, version number, GitHub derived activity measures, and the mean relative speed and accuracy rankings. Red colours indicate a positive correlation, blue colours indicate a negative correlation. Correlations with a P-value less than 0.05 (corrected for multiple-testing using the Benjamini-Hochberg method) are indicated with a ‘X’ symbol. The correlations with accuracy are illustrated in more detail in **B**, the relationship between speed and accuracy is shown in more detail in **Figure 2**. **B.** Violin plots of Spearman’s correlations for permuted accuracy ranks and different software features. The unpermuted correlations are indicated with a red asterisk. For each benchmark, 1,000 permuted sets of accuracy and speed ranks were generated, and the ranks were normalised to lie between 0 and 1 (see Methods for details). Circled asterisks are significant (empirical P-value < 0.05, corrected for multiple-testing using the Benjamini-Hochberg method).

Data on the number of updates to software tools from GitHub such as the version number, and numbers of contributors, commits, forks and issues was significantly correlated with software accuracy (respective Spearman’s rhos = 0.15, 0.21, 0.22, 0.23, 0.23 and respective Benjamini & Hochberg corrected P-values = 5.4 × 10^-4^, 1.1 × 10^-3^, 7.9 × 10^-4^, 3.4 × 10^-4^, 3.2 × 10^-4^, Additional file 1: Figures S6). The significance of these features was further confirmed with a permutation test (Figure 1B). These features were not correlated with speed however (see Figure 1A & Additional file 1: Figures S5&S6). We also found that reputation metrics such as citations, author and journal H-indices, and the age of tools were generally **not** correlated with either tool accuracy or speed (Figure 1A&B).

In order to gain a deeper understanding of the distribution of available bioinformatic software tools on a speed versus accuracy landscape, we ran a permutation test. The ranks extracted from each benchmark were randomly permuted, generating 1,000 randomized speed and accuracy ranks. In the cells of a 3 × 3 grid spanning the normalised speed and accuracy ranks we computed a Z-score for the observed number of tools in a cell, compared to the expected distributions generated by 1,000 randomized ranks. The results of this are shown in Figure 2. We identified 4 of 9 bins where there was a significant excess or dearth of tools. For example, there was an excess of “slow and inaccurate” software (Z=3.39, P-value=3.5 × 10^-4^), with more moderate excess of “slow and accurate” and “fast and accurate” software (Z=2.49 & 1.7, P=6.3 × 10^-3^ & 0.04 respectively). We find that only the “fast and inaccurate” extreme class is at approximately the expected proportions based upon the permutation test (Figure 2B).

**Figure 2:**
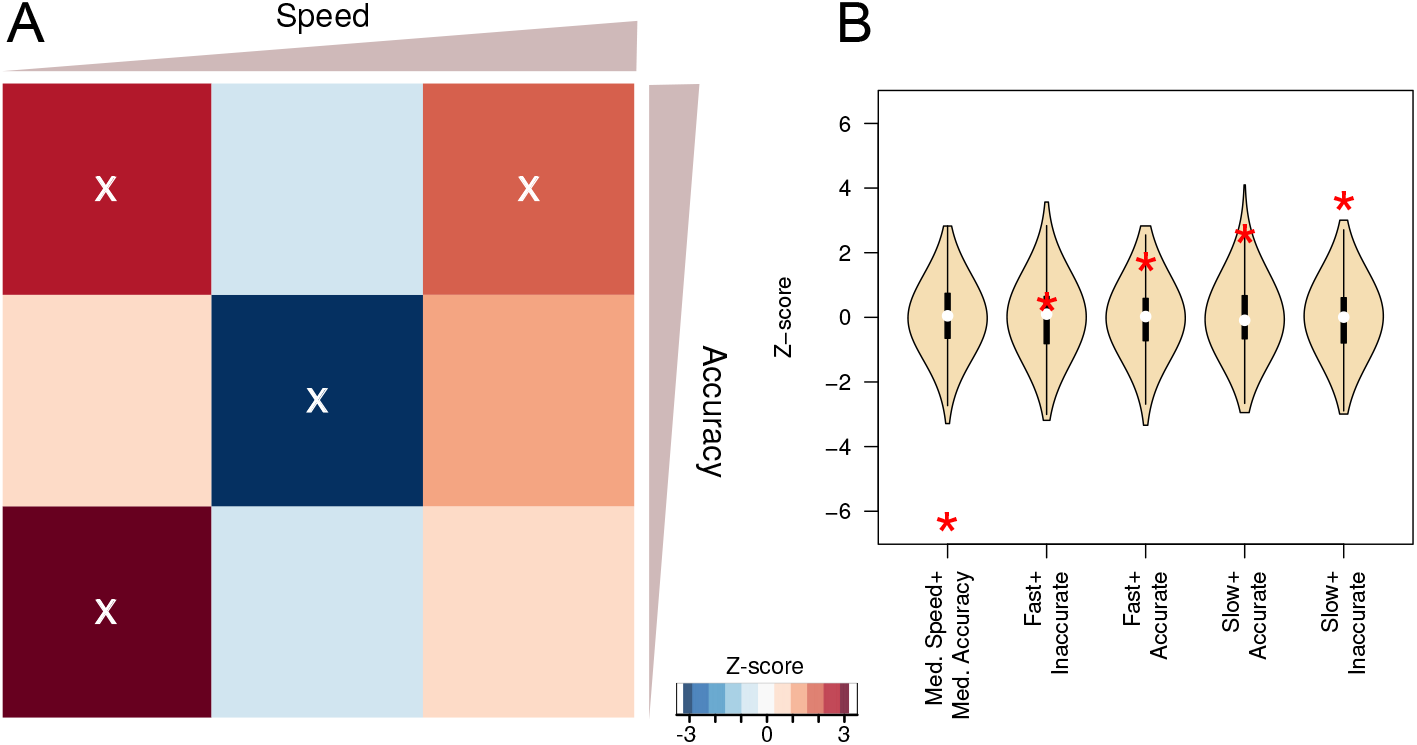
**A.** A heatmap indicating the relative paucity or abundance of software in the range of possible accuracy and speed rankings. Redder colours indicate an abundance of software tools in an accuracy and speed category, while bluer colours indicate scarcity of software in an accuracy and speed category. The abundance is quantified using a Z-score computation for each bin, this is derived from 1,000 random permutations of speed and accuracy ranks from each benchmark. Mean normalised ranks of accuracy and speed have been binned into 9 classes (a 3 × 3 grid) that range from comparatively slow and inaccurate to comparatively fast and accurate. Z-scores with a P-value less than 0.05 are indicated with a ‘X’. **B.** The z-score distributions from the permutation tests (indicated with the wheat coloured violin plots) compared to the z-score for the observed values for each of the corner and middle square of the heatmap.

The largest difference between the observed and expected software ranks is the reduction in the number of software tools that are classed as intermediate in terms of both speed and accuracy based on permutation tests (see Methods for details, Figure 2). The middle cell of Figure 2A and left-most violin plot of Figure 2B highlight this extreme, (Z=-6.38, P-value=9.0 × 10^-11^).

## Conclusion

We have gathered data on the relative speeds and accuracies of 499 bioinformatic tools from 69 benchmarks published between 2005 and 2020. Our results provide significant support for the suggestion that there are major benefits to the long-term support of software development [55]. The finding of a strong relationship between the number of commits and code contributors to GitHub (i.e. software updates) and accuracy, highlights the benefits of long-term or at least intensive development.

Our study finds little evidence to support that impact-based metrics have any relationship with software quality, which is unfortunate, as these are frequently cited reasons for selecting software tools [38]. This implies that high citation rates for bioinformatic software [1, 2, 3] is more a reflection of other factors such as user-friendliness or the Matthew Effect [56, 57] other than accuracy. Specifically, software tools published early are more likely to appear in high impact journals due to their perceived novelty and need. Yet without sustained maintenance these may be outperformed by subsequent tools, yet early publications still accrue citations from users, and all subsequent software publications as tools need to be compared in order to publish. Subsequent tools are not perceived to be as novel, hence appear in “lower” tier journals, despite being more reliable. Hence, the “rich” early publishers get richer in terms of citations. Indeed, citation counts are mainly predictive of age (Figure 1A).

We found the lack of a correlation between software speed and accuracy surprising. The slower software tools are over-represented at both high and low levels of accuracy, with older tools enriched in this group (Figure 2 and Additional file 1: Figure S7). In addition, there is an large under-representation of software that has intermediate levels of both accuracy and speed. A possible explanation for this is that bioinformatic software tools are bound by a form of publication-bias [58, 59]. That is, the probability that a study being published is influenced by the results it contains [60]. The community of developers, reviewers and editors may be unwilling to publish software that is not highly ranked on speed or accuracy. If correct, this may have unfortunate consequences as these tools may nevertheless have further uses.

While we have taken pains to mitigate many issues with our analysis, nevertheless some limitations remain. For example, it has proven difficult to verify if the gap in medium accuracy and medium speed software is genuinely the result of publication bias, or due to additional factors that we have not taken in to account. In addition, all of the features we have used here are moving targets. For example, as software tools are refined, their relative accuracies and speeds will change, the citation metrics, ages, and version control derived measures also change over time. Here we report a snapshot of values from 2020. The benchmarks themselves may also introduce biases into the study. For example, there are issues with a potential lack of independence between benchmarks (e.g., shared datasets, metrics and tools), there are heterogeneous measures of accuracy and speed and often unclear processes for including different tools.

We propose that the full spectrum of software tool accuracies and speeds serves a useful purpose to the research community. Like negative results, if honestly reported this information, illustrates to the research community that certain approaches are not practical research avenues [61]. The current novelty-seeking practices of many publishers, editors, reviewers and authors of software tools therefore may be depriving our community of tools for building effective and productive workflows. Indeed, the drive for novelty may be an actively harmful criteria for the software development community, just as it is for reliable and reproducible research [62]. Novelty-criteria for publication may, in addition, discourage continual, incremental improvements in code post-publication in favour of splashy new tools that are likely to accrue more citations.

In addition we suggest that further efforts be made to encourage continual updates to software tools. To paraphrase some of the suggestions of Siepel (2019), these efforts may include more secure positions for developers, institutional promotion criteria include software maintenance, lower publication barriers for significant software updates, encourage further funding for software maintenance and improvement - not just new tools [55]. If these issues were recognised by research managers, funders and reviewers, then perhaps the future bioinformatic software tool landscape will be much improved.

The most reliable way to identify accurate software tools is through neutral software benchmarks [54]. We are hopeful that this, along with steps to reduce the publication-bias we have described, will reduce the over-optimistic and misleading reporting of tool accuracy [26, 27, 29].

## Methods

In order to evaluate predictors of computational biology software accuracy, we mined the published literature, extracted data from articles, connected these with bibliometric databases, and tested for correlates with accuracy. We outline these steps in further detail below.

### Criteria for inclusion

We are interested in using computational biology benchmarks that satisfy Boulesteix’s (ALB) three criteria for a “neutral comparison study” [54]. Firstly, the main focus of the article is the comparison and **not** the introduction of a new tool as these can be biased [30]. Secondly, the authors should be reasonably neutral, which means that the authors should not generally have been involved in the development of the tools included in the benchmark. Thirdly, the test data and evaluation criteria should be sensible. This means that the test data should be independent of data that tools have been trained upon, and that the evaluation measures appropriately quantify correct and incorrect predictions. In addition, we excluded benchmarks with too few tools ≤ 3, or those where the results were inaccessible (no supplementary materials or poor figures).

### Literature mining

We identified an initial list of 10 benchmark articles that satisfy the ALB-criteria. These were identified based upon previous knowledge of published articles and were supplemented with several literature searches (e.g., [“benchmark” AND “cputime”] was used to query both GoogleScholar and Pubmed [53, 63]). We used these articles to seed a machine-learning approach for identifying further candidate articles and to identify new search terms to include. This is outlined in Additional file 1: Figure S1.

For our machine-learning-based literature screening, we computed a score, *s*(*a*), for each article that tells us the likelihood that it is a benchmark. In brief, our approaches uses 3 stages:

1. Remove high frequency words from the title and abstract of candidate articles (e.g. ‘the’, ‘and’, ‘of’, ‘to’, ‘a’, …)
2. Compute a log-odds score for the remaining words
3. Use a sum of log-odds scores to give a total score for candidate articles

For stage 1, we identified a list of high frequency (e.g. *f*(word) > 1/10, 000) words by pooling the content of two control texts [64, 65].

For stage 2, in order to compute a log-odds score for bioinformatic words, we computed the frequency of words that were not removed by our high frequency filter in two different groups of articles: bioinformatics-background and bioinformatics-benchmark articles. The text from bioinformatics-background articles were drawn from the bioinformatics literature, but these were not necessarily associated with benchmark studies. For background text we used Pubmed [53, 63] to select 8,908 articles that contained the word “bioinformatics” in the title or abstract and were published between 2013 and 2015. We computed frequencies for each word by combining text from titles and abstracts for the background and training articles. A log-odds score was computed for each word using the following formula:

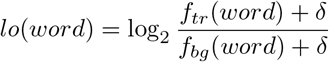

Where *δ* was a pseudo-count added for each word (*δ* = 10^-5^, by default), *f_bg_*(*word*) and *f_tr_*(*word*) were the frequencies of a *word* in the background and training datasets respectively. Word frequencies were computed by counting the number of times a word appears in the pool of titles and abstracts, the counts were normalised by the total number of words in each set. Additional file 1: Figure S2 shows exemplar word scores.

Thirdly, we also collected a group of candidate benchmark articles by mining Pubmed for articles that were likely to be benchmarks of bioinformatic software, these match the terms: “((bioinformatics) AND (algorithms OR programs OR software)) AND (accuracy OR assessment OR benchmark OR comparison OR performance) AND (speed OR time)”. Further terms used in this search were progressively added as relevant enriched terms were identified in later iterations. The final query is given in **Additional file 1**.

A score is computed for each candidate article by summing the log-odds scores for the words in title and abstract, i.e. 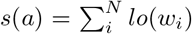. The high scoring candidate articles are then manually evaluated against the ALB-criteria. Accuracy and speed ranks were extracted from the articles that met the criteria, and these were added to the set of training articles. The evaluated candidate articles that did not meet the ALB-criteria were incorporated into the set of background articles. This process was iterated and resulted in the identification of 69benchmark articles, containing 134 different benchmarks. Together these ranked 499 distinct software packages.

There is a potential for bias to have been introduced into this dataset. Some possible forms of bias include converging on a niche group of benchmark studies due to the literature mining technique that we have used. A further possibility is that benchmark studies themselves are biased, either including very high performing or very low performing software tools. To address each of these concerns we have attempted to be as comprehensive as possible in terms of benchmark inclusion, as well as including comprehensive benchmarks (i.e., studies that include all available software tools that address a specific biological problem).

### Data extraction and processing

for each article that met the ALB-criteria and contained data on both the accuracy and speed from their tests, we extracted ranks for each tool. Until published datasets are made available in consistent, machine-readable formats this step is necessarily a manual process – ranks were extracted from a mixture of manuscript figures, tables and supplementary materials, each data source is documented in Additional file 2: Table S1. In addition, a variety of accuracy metrics are reported e.g. “accuracy”, “AUROC”, “F-measure”, “Gain”, “MCC”, “N50”, “PPV”, “precision”, “RMSD”, “sensitivity”, “TPR”, “tree error”, etc. Our analysis makes the necessarily pragmatic assumption that highly ranked tools on one accuracy metric will also be highly ranked on other accuracy metrics. Many articles contained multiple benchmarks, in these cases we recorded ranks from each of these, the provenance of which is stored with the accuracy metric and raw speed and accuracy rank data for each tool (Additional file 2: Table S1). In line with rank-based statistics, the cases where tools were tied were resolved by using a midpoint rank (e.g., if tools ranked 3 and 4 are tied, the rank 3.5 was used) [66]. Each rank extraction was independently verified by at least one other co-author to ensure both the provenance of the data could be established and that the ranks were correct. The ranks for each benchmark were then normalised to lie between 0 and 1 using the formula 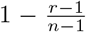 where ‘*r*’ is a tool’s rank and ‘*n*’ is the number of tools in the benchmark. For tools that were benchmarked multiple times with multiple metrics (e.g., BWA was evaluated in 6 different articles [67, 68, 69, 70, 71, 72]) a mean normalised rank was used to summarise the accuracy and speed performance. Or, more formally:

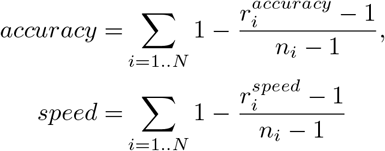

For each tool we identified the corresponding publications in GoogleScholar; the total number of citations was recorded, the corresponding authors were also identified, and if they had public GoogleScholar profiles, we extracted their H-index and calculated a M-index 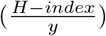 where ‘*y*’ is the number of years since their first publication. The journal quality was estimated using the H5-index from GoogleScholar Metrics.

The year of publication was also recorded for each tool. “Relative age” and “relative citations” were also computed for each tool. For each benchmark, software was ranked by year of first publication (or number of citations), ranks were assigned and then normalised as described above. Tools ranked in multiple evaluations were then assigned a mean value for “relative age” and “relative citations”.

The papers describing tools were checked for information on version numbers and links to GitHub. Google was also employed to identify GitHub repositories. When a repository was matched with a tool, the number of “commits” and number of “contributors” was collected, when details of version numbers were provided, these were also harvested. Version numbers are inconsistently used between groups, and may begin at either 0 or 1. To counter this issue we have added ‘1’ to all versions less than ‘1’, for example, version 0.31 become 1.31. In addition, multiple point releases may be used e.g. ‘version 5.2.6’, these have been mapped to the nearest decimal value ‘5.26’.

### Statistical analysis

For each tool we manually collected up to 12 different statistics from GoogleScholar, GitHub and directly from literature describing tools (1. corresponding author’s H-index, 2. corresponding author’s M-index, 3. journal H5 index, 4. normalised accuracy rank, 5. normalised speed rank, 6. number of citations, 7. relative age, 8. relative number of citations, 9. year first published, 10. version 11. number of commits to GitHub, 12. number of contributors to GitHub). These were evaluated in a pairwise fashion to produce Figure 1 A&B, the R code used to generate these is given in a GitHub repository (linked below).

For each benchmark of three or more tools, we extracted the published accuracy and speed ranks. In order to identify whether there was an enrichment of certain accuracy and speed pairings we constructed a permutation test. The individual accuracy and speed ranks were reassigned to tools in a random fashion and each new accuracy and speed rank pairing was recorded. For each benchmark this procedure was repeated 1,000 times. These permuted rankings were normalised and compared to the real rankings to produce the ‘X’ points in Figure 1B and the heatmap and histograms in Figure 2. The heatmap in Figure 2 is based upon Z-scores 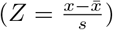. For each cell in a 3 × 3 grid a Z-score (and corresponding P-value is computed, either with the ‘pnorm’ distribution function in R (Figure 2A) or empirically (Figure 2B)) is computed to illustrate the abundance or lack of tools in a cell relative to the permuted data.

The distributions for each feature and permuted accuracy or speed ranks are shown in Additional file 1: Figures S3&S4. Scatter-plots for each pair of features is shown in Additional file 1: Figure S5. Plots showing the sample sizes for each tool, and feature are shown in Additional file 1: Figure S8, illustrates a power analysis to show what effect sizes we are likely to detect for our sample sizes.

## Supporting information

Supplementary figures S1-S8 and benchmark references.

## Acknowledgements

The authors acknowledge the valued contribution of invaluable discussions with Anne-Laure Boulesteix, Shinichi Nakagawa, Suetonia Palmer and Jason Tylianakis. Murray Cox, Raquel Norel, Alexandros Stamatakis, Jens Stoye, Tandy Warnow, and Luis Pedro Coelho and three anonymous reviewers provided valuable feedback on drafts of the manuscript.

This work was largely conducted on the traditional territory of Kāi Tahu.

## Availability of data and materials

Raw datasets, software and documents are available under a CC-BY license: https://github.com/Gardner-BinfLab/speed-vs-accuracy-meta-analysis

## Additional information

### Additional file 1

Supplementary Figures S1-S8 and the neutral software benchmark reference list the accuracy and speed data is derived from [1–66].

### Additional file 2

Supplementary Tables S1-S7.

