## Supplementary figures S1-S8 and benchmark references. for "Sustained software development, not number of citations or journal choice, is indicative of accurate bioinformatic software"

**Abstract**
In the below we provide additional results for our investigation of computational biology benchmarks.

**Keywords**
computational biology — accuracy — benchmarks — meta-analysis — software development

[1] *Department of Biochemistry, University of Otago, Dunedin, New Zealand.*
[2] *Biomolecular Interaction Centre, University of Canterbury, Christchurch, New Zealand.*
[3] *Department of Civil and Natural Resources Engineering, University of Canterbury, Christchurch, New Zealand.*
[4] *Parasites and Microbes, Wellcome Sanger Institute, Wellcome Genome Campus, Hinxton, 8 Cambridgeshire, CB10 1RQ, UK.*
[5] *Research Group for Genomic Epidemiology, National Food Institute, Technical University of Denmark, Kongens Lyngby, Denmark.*
[6] *Department of Research, Cancer Registry of Norway, Oslo, Norway.*
[7] *New Zealand eScience Infrastructure, 49 Symonds St, Auckland, New Zealand.*
[8] *Department of Computer Science, University of Otago, Dunedin, New Zealand.*

### Literature mining

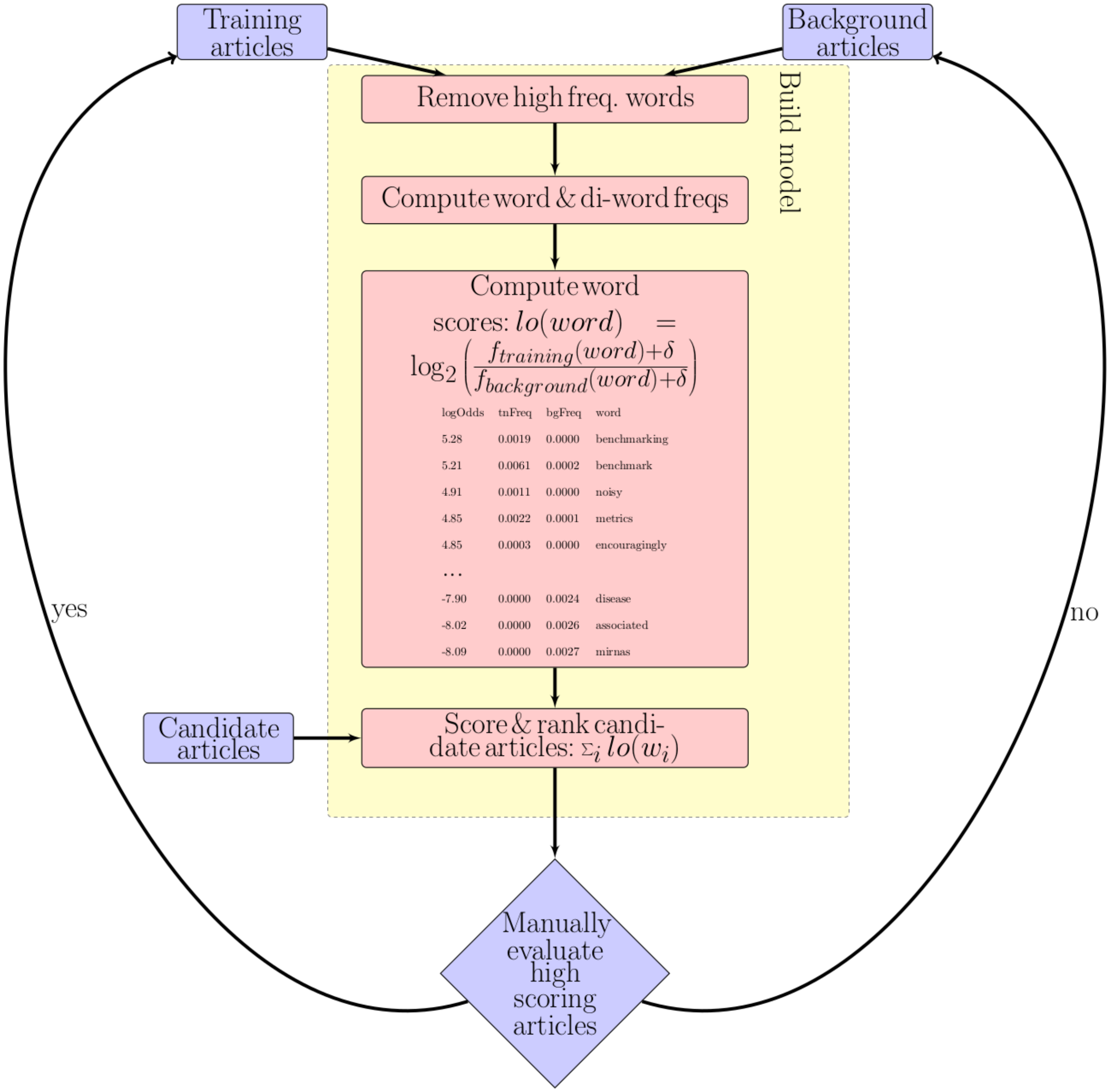


**Figure S1.** In order to improve the identification of benchmark articles that rank both accuracy and speed we developed a tool for ranking PubMed articles based upon word association scores (measured in ‘bits’). In brief, keywords were extracted from titles and abstracts for both training (in this case benchmark articles) and background articles (articles published between 2013 and 2015 with “bioinformatics” in the title or abstract). Log-odds ratios were computed for each keyword (measured in ‘bits’). Candidate articles that matched a hand-selected list of keywords associated with benchmarks: `((bioinformatics OR (computational AND biology)) AND (algorithmic OR algorithms OR biotechnologies OR computational OR kernel OR methods OR procedure OR programs OR software OR technologies)) AND (accuracy OR analysis OR assessment OR benchmark OR benchmarking OR biases OR comparing OR comparison OR comparisons OR comparative OR comprehensive OR effectiveness OR estimation OR evaluation OR metrics OR efficiency OR performance OR perspective OR quality OR rated OR robust OR strengths OR suitable OR suitability OR superior OR survey OR weaknesses OR correctness OR correct OR evaluate OR competing OR competition) AND (complexity OR cputime OR runtime OR walltime OR duration OR elapsed OR fast OR faster OR perform OR performance OR slow OR slower OR speed OR time OR (computational AND resources))`. These were then scored and ranked with a “sum of bits” score. High ranking articles were then inspected, those that met our criteria were added to the training set, those that did not were added to the background set of articles.

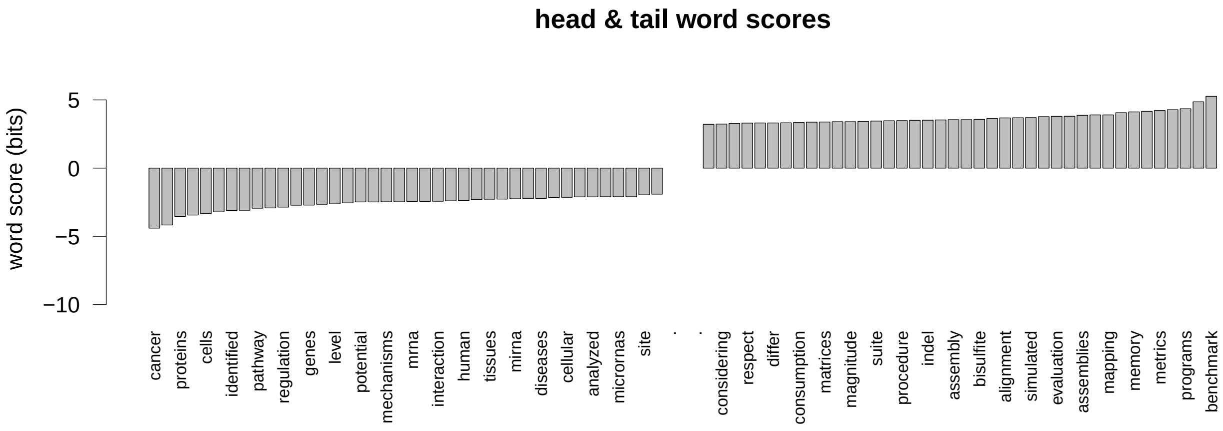


**Figure S2.** The 40 highest (**right**) and lowest (**left**) scoring words that are associated with bioinformatic benchmark articles from the training benchmark articles, compared to the background articles. The log-odds ratios (measured in bits) are indicated on the y-axis.

### Data visualisations

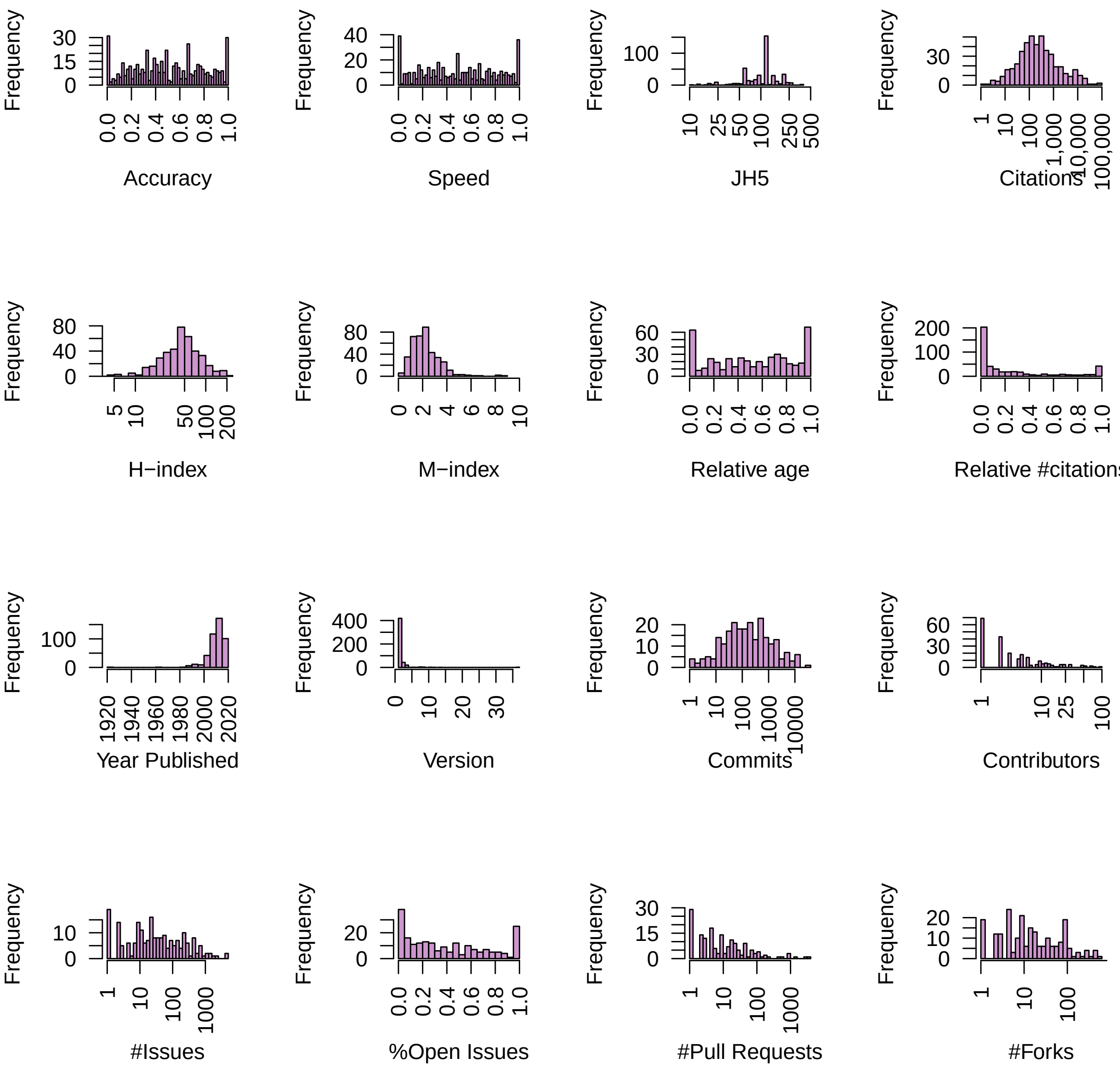


**Figure S3.** The distributions for the measures that have been proposed to be predictive of software quality used in this study. These are, reading from left to right, top to bottom: Accuracy – the mean normalised accuracy rank for each benchmarked method; Speed – the mean normalised speed rank for each benchmarked method; JH5 – the Journal H5 index to the highest impact journal that has published a manuscript describing a method, data from GoogleScholar 2020 Metrics; Cites – the number of citations to the most cited manuscript describing a method, data from GoogleScholar; H-index – the H-index for the highest profile corresponding author from the manuscripts describing a method, data from GoogleScholar User Profiles; M-index – the M-index (H-index/(#years since first publication)) for the highest profile corresponding author from the manuscripts describing a method, data from GoogleScholar User Profiles; Relative ages – for different tools, for each benchmark tools were ranked based upon publication dates, these ranks were normalised to lie between 0 and 1. Relative #citations – within each benchmark, what is the relative age of each tool, this corrects for older research fields (e.g. sequence alignment) vs newer research fields (e.g. single-cell sequencing); Year published – the first year each software tool was published; Version – the most resent version number for each tool; Commits – the number of commits to Github; Contributors – the number of contributors to a Github repository; #Issues – the number of open and closed issues for each Github repository; %Open issues – the pecentage of issues that remain open for each Github repository; #Pull requests – the number of pull requests to a Github repository; #Forks – the number of times a Github repository has been forked;

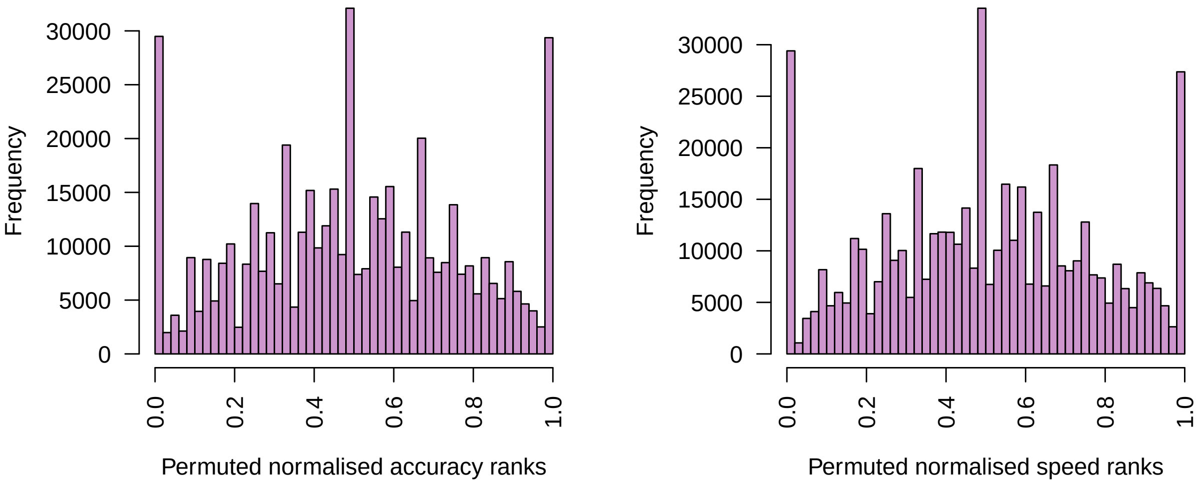


**Figure S4.** The distributions for the permuted accuracy (**left**) and speed (**right**) values, where accuracy and speed ranks have been normalised to lie between 0 and 1.

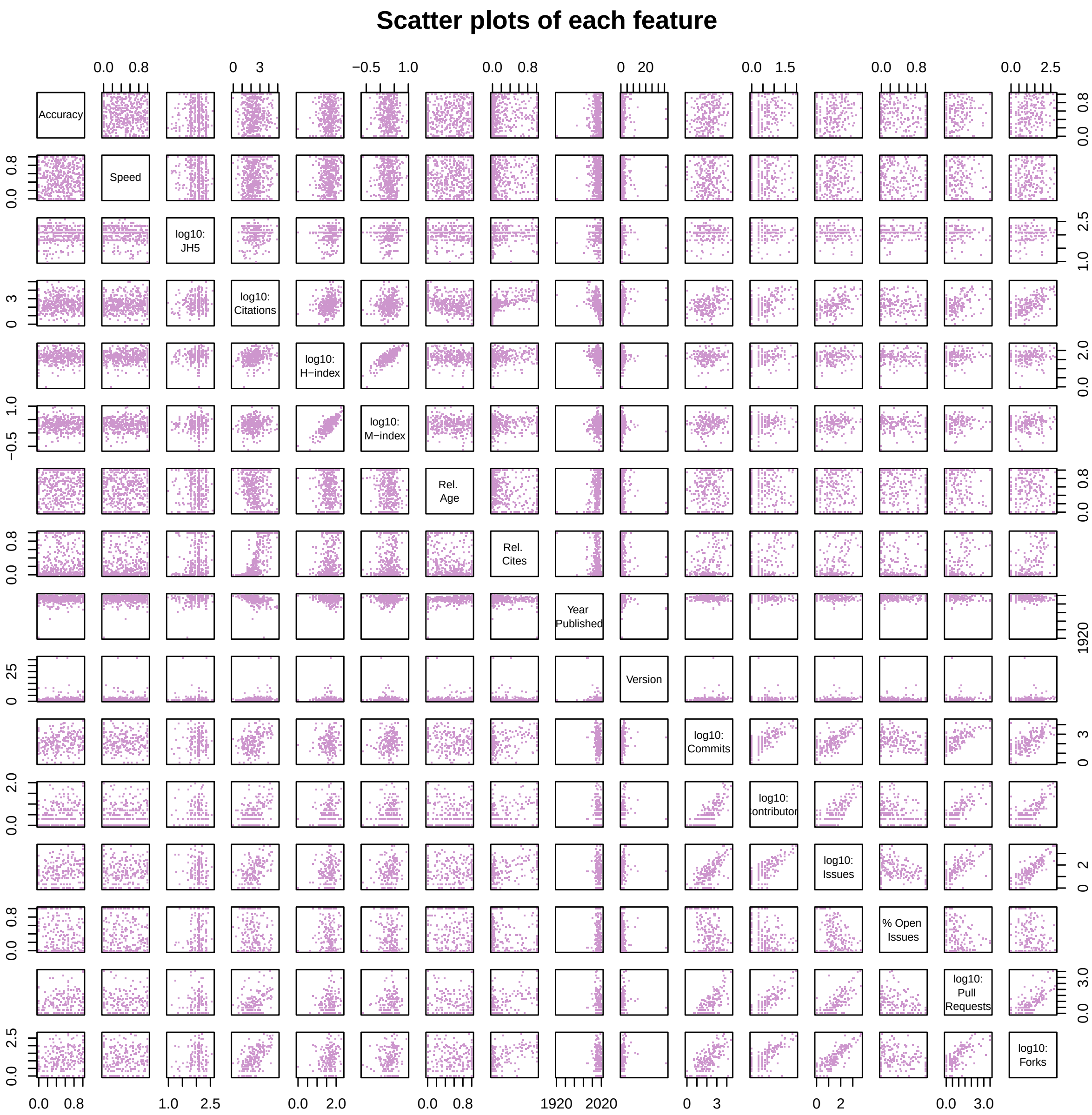


**Figure S5.** Dot plots for each feature relative to every other feature (accuracy, speed, JH5, citations, H-index, M-index, relAge, relCites, yearPublished, version, commits, contributors, #Issues, %Open issues, #Pull requests, and the #Forks).

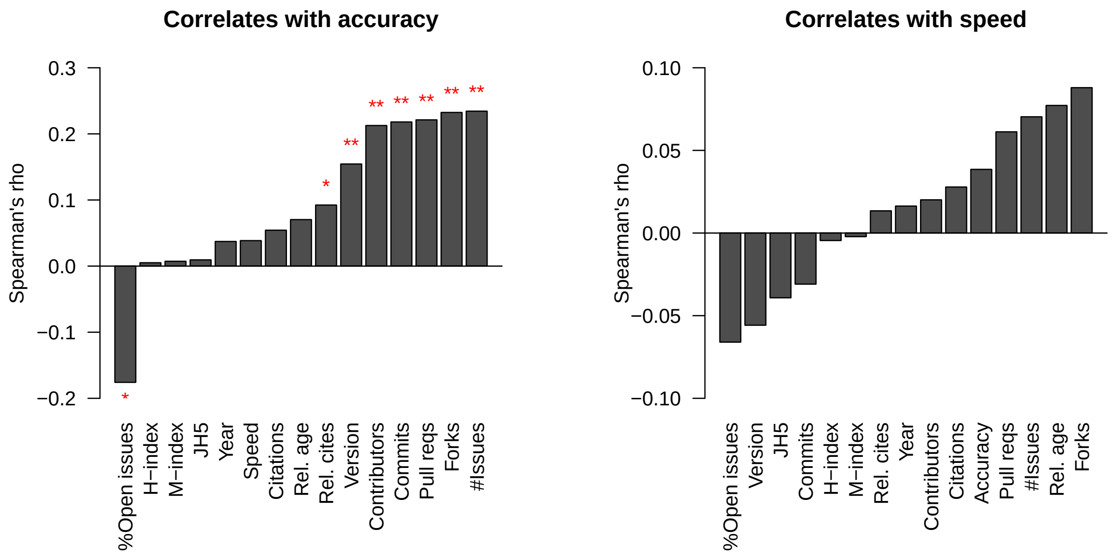


**Figure S6.** The correlation between method accuracy (**left**) and method speed (**right**) and measures we expected to be predictive of software quality. E.g. author reputation measures (H-index, M-index), journal reputation (JH5 and JIF), number of users (citations and relative citations) or the recency of methods (year and relative age). The correlations were estimated using Spearman's $\rho$. The signficant relationships are indicated with a "*".

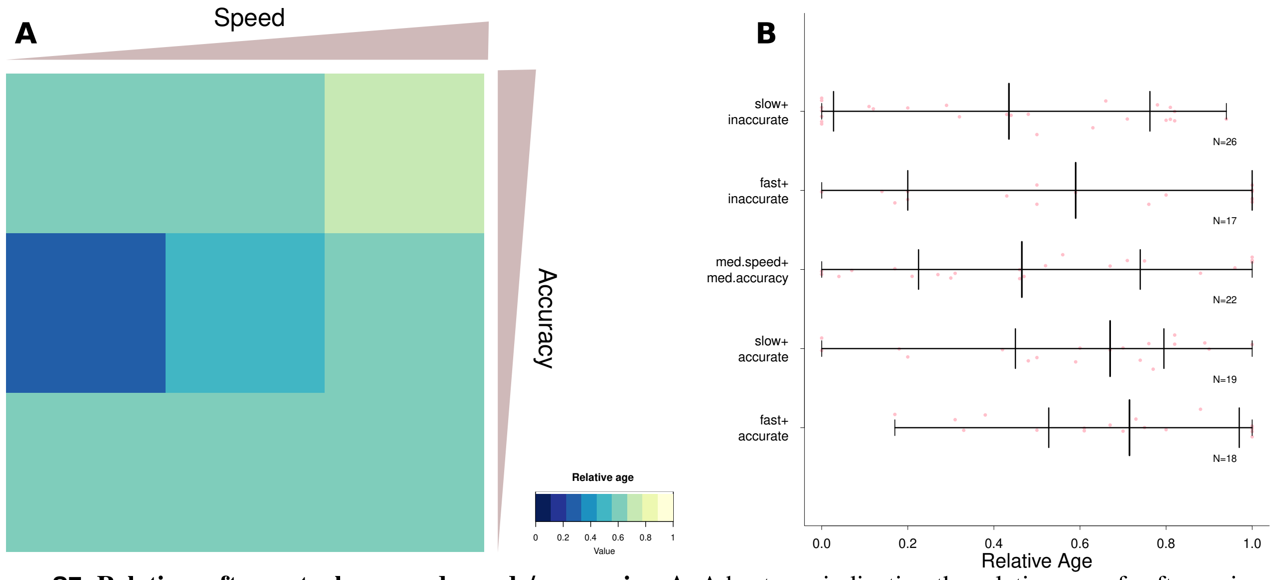


**Figure S7. Relative software tool ages and speeds/accuracies. A.** A heatmap indicating the relative age of software in a range of relative accuracy and speed rankings. Blue colours indicate an abundance of older software tools in an accuracy and speed category, while light colours indicate younger software in an accuracy and speed category. **B.** Jitter-plots for the relative age distribution for software tools in each of the 5 3x3 cells indicated in **B**. The five row correspond to the four extreme corners of the speed vs accuracy spectrum (i.e. slow and inaccurate, slow and accurate, fast and inaccurate, fast and accurate) and the central box (medium speed and medium accuracy). Median, quartile and extreme values are indicated with tick marks for each group. Accurate and fast software tools tend to have been published more recently than inaccurate and slow tools (Wilcox rank sum test, $W = 341,\ P = 5 \times 10^{-3}$)

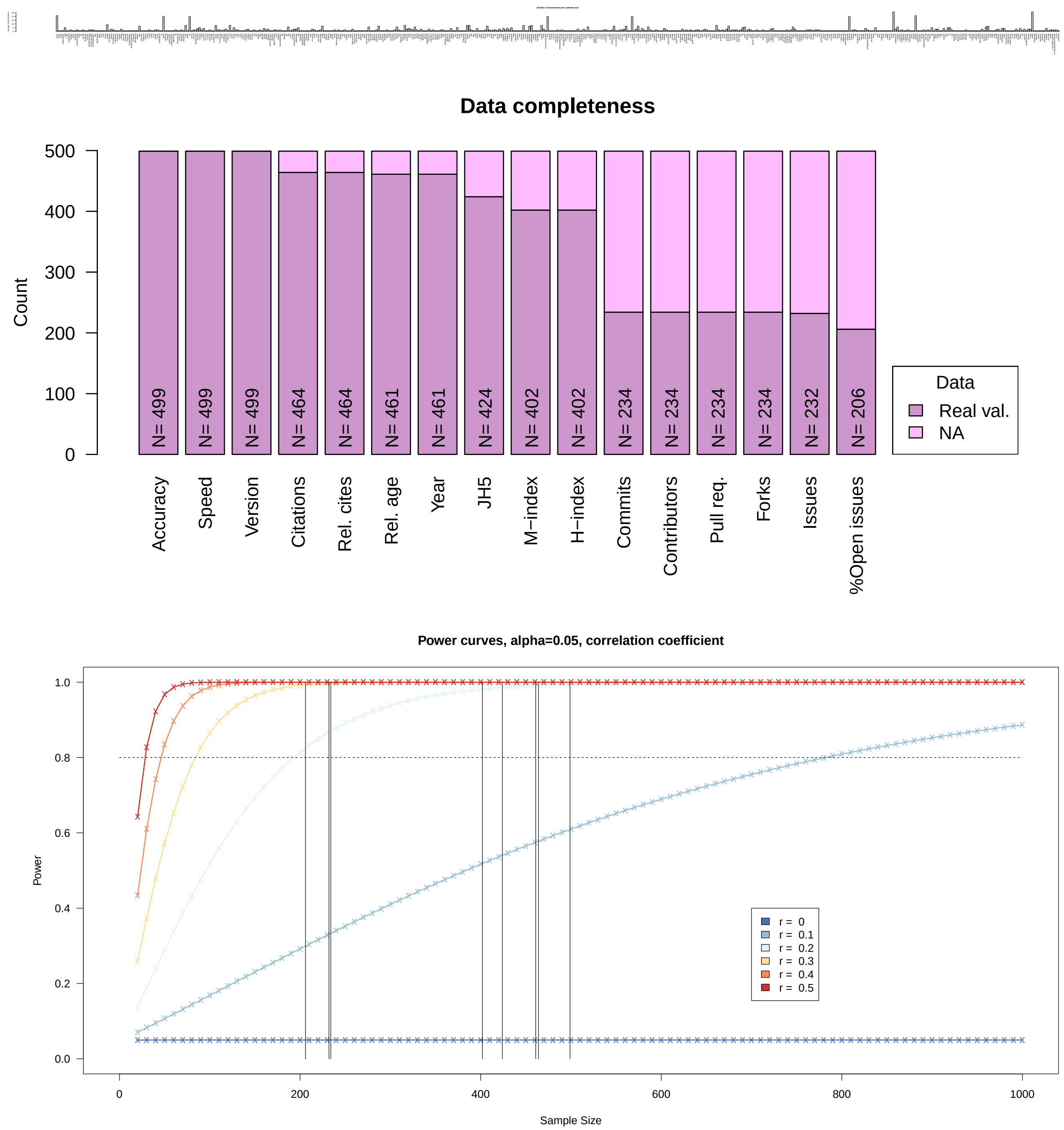


**Figure S8. Top:** The number of benchmarks each software tool has appeared in. **Middle:** Plot of the amount missing/known data for each of the software features used in this analysis. **Bottom:** a power curve analysis shows the sample sizes for each set of features is sufficient to detect correlations $r \geq 0.2$ between our features of interest with $power \geq 0.8$. For example, for speed vs accuracy $N = 499$, which is indicated with a vertical line.

The complete list of benchmarks used for this study [1, 2, 3, 4, 5, 6, 7, 8, 9, 10, 11, 12, 13, 14, 15, 16, 17, 18, 19, 20, 21, 22, 23, 24, 25, 26, 27, 28, 29, 30, 31, 32, 33, 34, 35, 36, 37, 38, 39, 40, 41, 42, 43, 44, 45, 46, 47, 48, 49, 50, 51, 52, 53, 54, 55, 56, 57, 58, 59, 60, 61, 62, 63, 64, 65, 66].
